## Supplementary Material for "Linear-regression-based algorithms can succeed at identifying microbial functional groups despite the nonlinearity of ecological function"

#### 1 Derivation of Eq. (3) in main text

Substituting $D_{ij} = \delta_{i,j+1}$ and $R_i = R_1\delta_{1i}$ into Eq. (1b), one obtains

$$\begin{aligned}
\frac{dm_1}{dt} &= R_1 - m_1 T_1 - d_1 m_1,\\
\frac{dm_2}{dt} &= w_1 T_1 m_1 - m_2 T_2 - d_2 m_2,\\
&\cdots\\
\frac{dm_{N-1}}{dt} &= w_{N-2} T_{N-2} m_{N-2} - m_{N-1} T_{N-1} - d_{N-1} m_{N-1},\\
\frac{dm_N}{dt} &= w_{N-1} T_{N-1} m_{N-1} - d_N m_N,
\end{aligned}$$

where $T_i = \sum_\mu \tau_{i\mu} n_\mu$ is the total abundance of functional group $i$ ($i \in \{1, \ldots, N-1\}$). Equating the left hand side to 0, one obtains successively

$$\begin{aligned}
m_1 &= \frac{R_1}{T_1 + d_1},\\
m_2 &= \frac{w_1 T_1 m_1}{T_2 + d_2} = R_1 \frac{w_1 T_1}{T_1 + d_1} \frac{1}{T_2 + d_2},\\
&\cdots\\
m_{N-1} &= R_1 \frac{w_1 T_1}{T_1 + d_1} \frac{w_2 T_2}{T_2 + d_2} \cdots \frac{1}{T_{N-1} + d_N},\\
m_N &= \frac{w_{N-1} T_{N-1} m_{N-1}}{d_N} = \frac{R_1}{k_N} \frac{w_1 T_1}{T_1 + d_1} \frac{w_2 T_2}{T_2 + d_2} \cdots \frac{w_N - 1 T_{N-1}}{T_{N-1} + d_N},
\end{aligned}$$

with the last line being Eq. (3) in the main text.

#### 2 Recovery quality, grouping score and performance ceiling of a k-group grouping under N-group ground truth

We denote the set of all species as $\mathcal{S} = \{s_1, \ldots, s_S\}$. A $k$-group grouping of all species is a set $P_k = \{g_1, \ldots, g_k\}$ satisfying

$$g_i \cap g_j = 0 \;\forall g_i, g_j \in P_k, i \neq j \tag{S1a}$$

$$\bigcup_{i=1}^{k} g_i = \mathcal{S}, \tag{S1b}$$

where $g_i = \{s_{i_1}, \ldots, s_{i_m}\}$ is a group of $m$ species ($m$ can be different for each group).

Let's denote the ground truth grouping as $\mathcal{T}_N = \{t_1, \ldots, t_N\}$ which contain $N$ disjoint groups, and let $P_k$ be a $k$-group grouping returned by one of our algorithms, $P_k = \{g_1, \ldots, g_k\}$. For each pair of $t_i \in \mathcal{T}_N$ and $g_j \in P_k$, we calculate the Jaccard Similarity $J_{ij}$ between them, which is defined as

$$J_{ij} = \frac{|t_i \cap g_j|}{|t_i \cup g_j|}, \tag{S2}$$

where $|a|$ means number of elements in the set $a$. The largest $J_{ij}$ for a given $i$ is defined as the *recovery quality* of a true group $t_i$ by the grouping $P_k$, and the average of the recovery qualities of all $t_i$ is defined to be the overall quality *score* of the grouping:

$$\text{Recover}_{\,t_i}(P_k) \equiv \max_j (J_{ij}) \tag{S3a}$$

$$\text{Score}_{\,\mathcal{T}_N}(P_k) \equiv \frac{1}{N}\sum_{i=1}^{N} \text{Recover}_{\,t_i}(P_k). \tag{S3b}$$

Given an $N$-group ground truth $\mathcal{T}_N$ and an integer $k \leq N$, the quality score of a $k$-group grouping $P_k$ will be highest if and only if every group $g_j \in P_k$ is either identical to one of the true groups $t_i \in \mathcal{T}_\mathcal{N}$, or is the union of several true groups. In other words, a $k$-group grouping $P_k$ will score highest if the indices of the true groups $1 \ldots N$ can be partitioned into $k$ non-overlapping subsets $G_i$ ($i \in 1 \ldots k$), such that:

$$G_i \cap G_j = 0 \;\forall i \neq j \tag{S4a}$$

$$\bigcup_{j=1}^{k} G_j = \{1, 2 \ldots N\} \tag{S4b}$$

$$\forall g_j \in P_k : g_j = \bigcup_{i \in G_j} t_i \tag{S4c}$$

The overall quality score of $P_k$ is the average recovery quality of each group, and this average can now be easily computed by grouping the summation indices $1 \ldots N$ according to $G_i$:

$$\text{Score}_{\,\mathcal{T}_N}(P_k) \equiv \frac{1}{N}\sum_{i=1}^{N} \text{Recover}_{\,t_i}(P_k) = \frac{1}{N}\sum_{j=1}^{k}\left[\sum_{i \in G_j} \text{Recover}_{\,t_i}(P_k)\right], \tag{S5}$$

and in this expression, the term in the square bracket simplifies to 1 (independently of $j$):

$$\sum_{i \in G_j} \text{Recover}_{\,t_i}(P_k) \equiv \sum_{i \in G_j} \text{Jaccard}\,(t_i, g_j) = \sum_{i \in G_j} \frac{|t_i \cap g_j|}{|t_i \cup g_j|} = \frac{\sum_{i \in G_j} |t_i|}{|g_j|} = \frac{|g_j|}{|g_j|} = 1, \tag{S6}$$

Here, we used the fact that in this case, the recovery quality of a true group $t_i \in G_j$ is the Jaccard similarity between $t_i$ and the group in $P_k$ that contains it. Thus, we conclude that for $N$-group ground truth, the *performance ceiling* of a $k$-group grouping (with $k \leq N$) is

$$\text{Score}_{\,\mathcal{T}_N}(P_k) = \frac{1}{N}\sum_{j=1}^{k}\left[\sum_{i \in G_j} \text{Recover}_{\,t_i}(P_k)\right] = \frac{k}{N}. \tag{S7}$$

Fig. S1 shows some examples of 2-group, 3-group and 4-group groupings under a 3-group ground truth with their scores shown on the right. The figure illustrates several features of our scoring metric. First, groupings 2-4 all achieve score 2/3, which is the performance ceiling of a 2-group grouping given a 3-group ground truth. Second, sometimes a $k$-group grouping returned by an algorithm is essentially a $(k-1)$-group one, e.g., groupings 1 and 7, which is appropriately reflected in their scores. Third, as expected, errors will lower the score (and score penalty increases with the number of errors), as seen by comparing groupings 5 and 6, or groupings 9 and 10. These points support our definition as being a good scoring metric. Finally, the figure also illustrates a limitation of our definition when $k > N$. For the two 4-group groupings shown (11 & 12), our score clearly favors grouping 11, but it is arguably not obvious which of these two candidate groupings should be treated as "more correct".

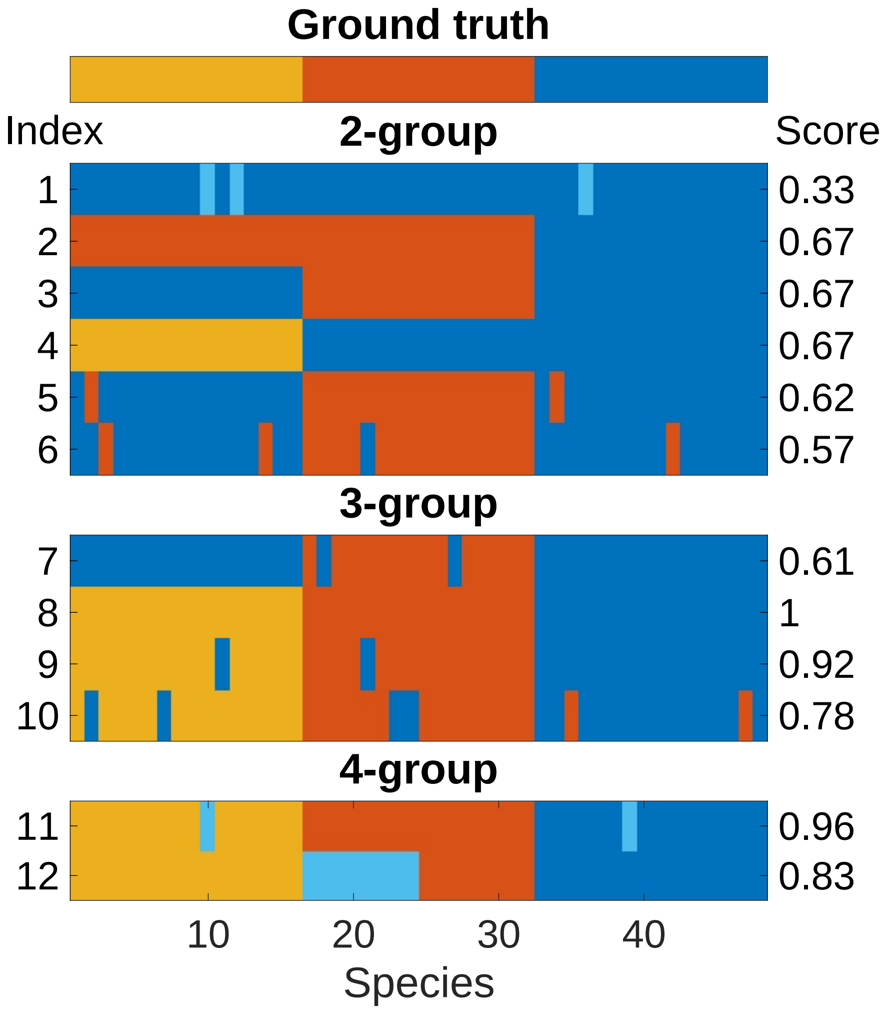


Figure S1: Example 2-group, 3-group and 4-group groupings under a 3-group ground truth with their scores shown on the right.

#### 3 Effect of "nonfunctional group" on function

In main text we mentioned that while EQO groups species into two groups, it assumes that only one of them (the "functional group") affects the level of function, i.e., it assumes that the remaining species (the "nonfunctional group", quotes will be omitted later for simplicity) have no effect on function. However, here we show that, at least in our model, the nonfunctional group will in general have negative effect on the level of function due to competition with the functional species. Consequently, at least in our synthetic datasets, the assumption built into EQO will underestimate the predictive power of a grouping, thus explaining the comparatively low performance score of EQO in Fig. 2 of the main text.

To exclude the complexity introduced by multi-functional groups, here we focus on the case where there's only one functional group. We let $N = 2$ in the model so there's only one step in the degradation chain which can be catalyzed by the functional group. As in Fig. 3 of the main text we keep the total number of species to be 48, with the functional and nonfunctional group both containing 24 species, and set the external supply of metabolite 1 to $R_1 = w^{-(N-1)} = 2$.

In this model (and probably in many real communities) species in the nonfunctional group compete with those in the functional one. In our model, this competition is implemented by the generalised depletable resources $h_a$, held fixed. Thus, the abundances of the 2 groups will be anti-correlated with each other. Since the functional group will positively correlate with function, we expect the nonfunctional group to have an indirect anti-correlation with the function. Consequently, in our model the non-functional species will in general have negative coefficients under a regression of function against species. Both these expectations are confirmed in Fig. S2A & B, where for one example dataset we plot the function against the abundance of nonfunctional group in panel A, which shows a negative correlation; and show the coefficients of all species for a linear regression against function in panel B, where we see all nonfunctional species (blue) have negative coefficients. The clear separation of coefficients of species in the two groups also explains the success of the simple K-Means algorithm .

As described in Materials and methods of the main text, our implementation of EQO is equivalent to performing a 1-dimensional regression of function against the abundance of the putative functional group. We note that this regression can also be viewed as an $S$-dimensional regression of function against all $S$ species with the restriction that coefficients of species in the functional group equal to a same nonzero value, while the coefficients of species in the nonfunctional group are constrained to equal 0. From Fig. S2B we find that the first restriction can be a good approximation (since the coefficients of functional species are similar), while the second can be slightly worse, since the mean coefficient of nonfunctional species is negative and is of the same order as that of functional species (i.e., not negligible). Thus, we hypothesized that this zero-coefficient restriction on nonfunctional group will hamper the performance of EQO.

To test this, we introduce a variant of EQO where both group can have nonzero coefficients, which is termed "EQO-2g", with the "2g" suffix indicating that both groups are included in the regression. As in EQO (see Materials and methods in the main text), we use a Boolean vector $\vec{x} \equiv \{x_\mu\}$ of length $S$ to represent a grouping of all $S$ species, which we seek to optimize. For a given $\vec{x}$, we calculate both the abundance of the functional group (the species corresponding to $x_\mu = 1$) and the abundance of the nonfunctional group (the species corresponding to $x_\mu = 0$). We then perform a two-dimensional linear regression of function $Y_a$ against the abundances of the two groups (with an intercept). In this variant, the number of parameters in the model does not change with the sizes of the two groups. Thus, there is no need for the AIC as the model-selection criterion, and we can directly use RSS (residual sum of squares) as the objective function to minimize. As in EQO we use the MATLAB's genetic-algorithm optimizer `ga` (with the same options mentioned in the EQO section of Materials and methods in main text). We note that this version of EQO is substantially equivalent to the 2-group Metropolis, except the optimization is performed by genetic algorithm (as in the original EQO paper), rather than the Metropolis-like procedure described in Materials and methods in main text. Thus, the performance of EQO-2g can be seen as a control for evaluating the perforamnce of both EQO and Metropolis.

As shown in Fig. S2C, EQO-2g performs as well as K-Means and Metropolis, and all three slightly outperform EQO, as before.

As a final test, we doubled the number of general resources (from 15 to 30). Our expectation is that this should decrease the competition-mediated effect of the nonfunctional group on the functional species, and thus on the function — conditions which may favour EQO performance. The results are shown in Fig. S2D. We see that the performance of EQO improves a lot, as expected. Putting these observations together, we conclude that allowing the species not directly involved in the function to have nonzero coefficients is likely beneficial to the performance of regression-based group-searching algorithms.

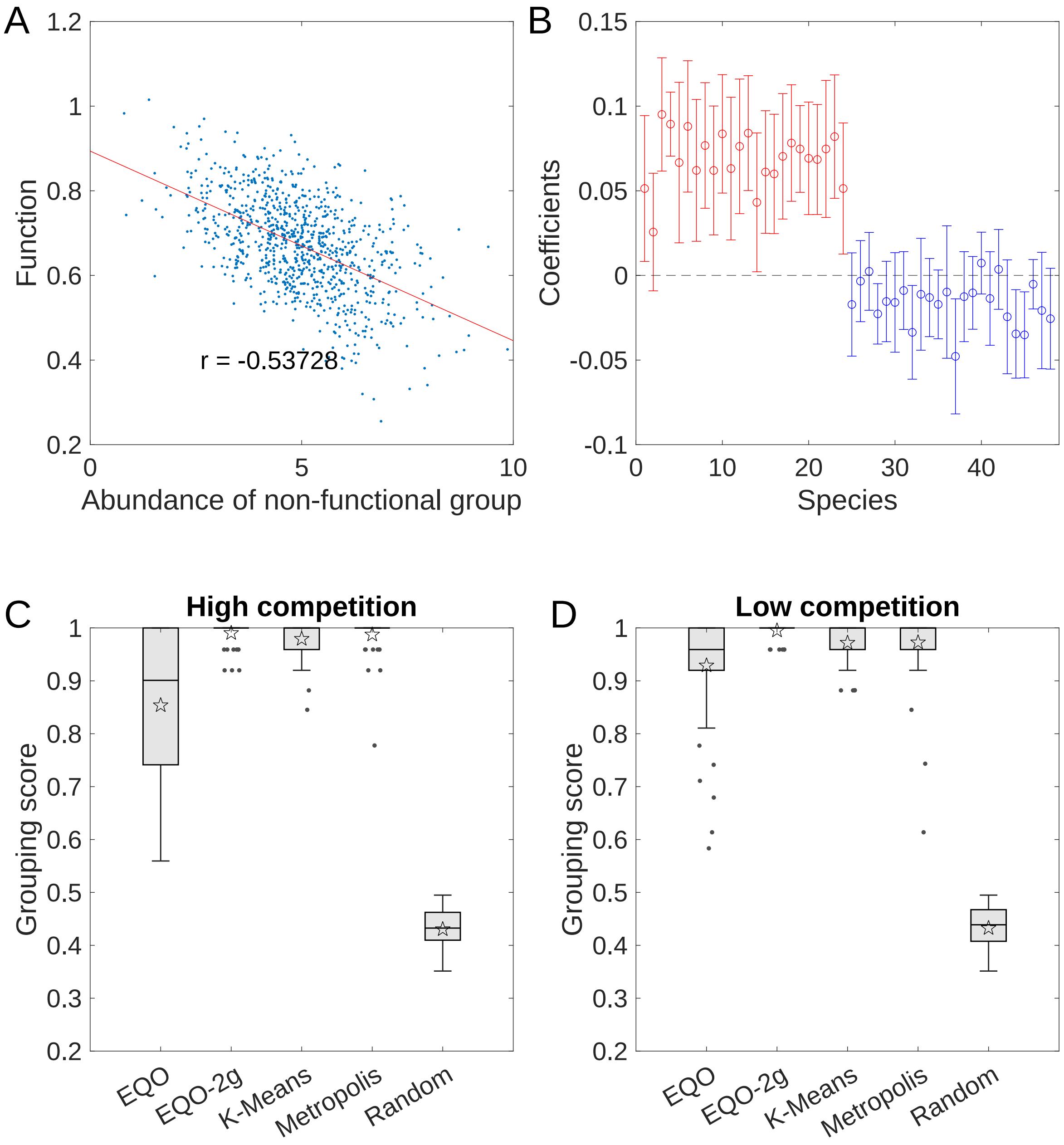


Figure S2: **Effect of "nonfunctional group" on function.** We consider the model in main text with $N = 2$. (A) The value of function (the final metabolite) shown against the abundance of the nonfunctional group (species not involved in producing this metabolite) for an example dataset of 900 samples. The scatter plot shows a negative correlation. The red line is the least-squares line and r marks the Pearson correlation. (B) The coefficients of all species of a $S$–dimensional regression of function against all $S$ species, for the same dataset as in (A). Error bars indicate 95% confidence intervals for the coefficient estimates. The $x$ axis is ordered so that species 1-24 belong to the functional group (red) and species 25-48 belong to the nonfunctional group (blue). We see that most nonfunctional species have negative regression coefficients. (C) The grouping scores for the outputs of EQO, EQO-2g, K-Means and Metropolis algorithms over 50 simulated datasets, shown as box plot with markers the same as Fig. 2A&B. EQO-2g performs as well as Metropolis. Random groupings are included as controls. (D) Same as (C), with the competition strength tuned down by doubling the number of general resources. The performance of EQO is improved and comparable to other algorithms, as expected (see text).

### 4 Performance of linear-regression-based algorithms in other model ecological scenarios

Here we test the generality of our results in the main text by testing the algorithms on 3 other functions. For clarity we only show results for the Metropolis algorithm.

Specifically, we let the function to be (1) an intermediate (rather than the final) product of a linear degradation chain; (2) one of the end products in a branched degradation chain; and (3) the common end product of 2 convergent degradation chains. We find that, in the first two cases, Metropolis can identify all the functional groups, while in the third, it can only recover the group of direct producers of the metabolite of interest. Put together,for a function associated with multiple groups, in general the difficulty for finding different groups will be different (due to their different positions in the metabolism structure), and the group which affects (is correlated with) the function the most will be easiest to recover (and sometimes the *only* recoverable one).

#### 4.1 An intermediate product of a linear degradation chain

This model is identical to the linear degradation chain model in the main text with $N = 3$ metabolites; the only difference is that the function of interest is now defined as the concentration of the intermediate degradation product (metabolite 2). We find that the Metropolis algorithm behaves similar to the scenario discussed in the main text (when the function of interest is the concentration of the end product). As shown in Fig. S3A, the 2-group output detects group 2, whereas the 3-group output additionally resolves the species belonging to group 1.

#### 4.2 Branched metabolic structures

The two other metabolic strctures we consider are a degradation chain with a branch and two linear degradation chains sharing the same source and end product. One important difference between these new structures and the single linear chain discussed above is that now a metabolite can serve as a substrate of more than one reactions. To accommodate this, we modify our model equations as follows:

$$\frac{dn_\mu}{dt} = n_\mu \left( \sum_r (1 - w_r)\, \tau_{r\mu} m_{i_r} + \sum_a \sigma_{a\mu} h_a - \chi_\mu \right), \tag{S8a}$$

$$\frac{dm_i}{dt} = R_i + \sum_{r,\mu} C_{ir} n_\mu \tau_{r\mu} m_{i_r} - d_i m_i, \tag{S8b}$$

$$h_a = h_a\left(\{\sigma_{a\nu}\}, \{n_\nu\}\right) = \frac{h_0^a}{1 + \sum_\nu \sigma_{a\nu} n_\nu / K_a}. \tag{S8c}$$

Now, the matrix $\{\tau_{r\mu}\}$ describes which species catalyze which *reactions*, enumerated by the index $r$. ($\tau_{r\mu}$ equals 1 is species $\mu$ can catalyze reaction $r$.) As before, for the groups to be clearly defined, we require that each species can catalyze at most one reaction. $\{C_{ir}\}$ is the stoichiometric matrix with its $r$'th column corresponding to reaction $r$. Specifically, if reaction $r$ transfers a fraction $w_r$ of metabolite $i$ into $j$, then $C_{ir} = -1$, $C_{jr} = w_r$, and $C_{kr} = 0$ for $k \neq i, j$. We denote $i_r$ to be the label of the substrate of reaction $r$ (i.e., $C_{i_r r} \equiv -1$). Other symbols are the same as Eq. (1) in the main text. External supply is set to $R_i = R_1 \delta_{1i}$, with $R_1$ fixed to be 4 for both two models. Other parameters are the same as in the main text, unless stated otherwise.

##### 4.2.1 A degradation chain with a branch

In this model we have 3 reactions (Fig. S3B), and we let each reaction to be catalyzed by 12 species, which form 3 non-overlapping functional groups. In addition, we include 12 species not involved in the degradation; the total number is therefore $S = 48$ species as before. As in the case of linear degradation chain, the group of direct producers (group 2) is still the one recovered best, and the only one identified in 2-group outputs. What's more, remarkably, all 3 functional groups can be recovered in the 4-group outputs.

##### 4.2.2 Two linear degradation chains sharing the same source and end product

In this model, there are two alternative pathways from the source metabolite (#1) to the end-product metabolite (#4). To distinguish them, we let the transfer ratio $w_r$ to take different values for different reactions as indicated in Fig. S3C, left. (As a reminder, in the other models all transfer ratios were set to a fixed value of 0.5.) There are 8 species to catalyze each of the 4 reactions, together with 16 other species not involved in the function ($S = 48$ species in total). At a finer resolution, the functional species can be divided into 4 groups, each for one reaction, as indicated in Fig. S3C, left. At a coarser resolution, however, the two groups catalyzing the the 1st reactions of the two pathways (group 1a and 1b) can be combined into a single "group 1", while the two groups of direct producers (group 2a and 2b) can be combined into "group 2". Interestingly, we see that if we ask for a 2-group grouping, the algorithm distinguishes all direct producers (group 2) from the rest (Fig. S3C, middle), while if we ask for a 3-group grouping, rather than identifying upstream species it will subdivide group 2 into group 2a and 2b (Fig. S3C, right). Further increasing number of groups does not lead to the recovery of upstream group(s). Thus, in this scenario only direct producers can be found.

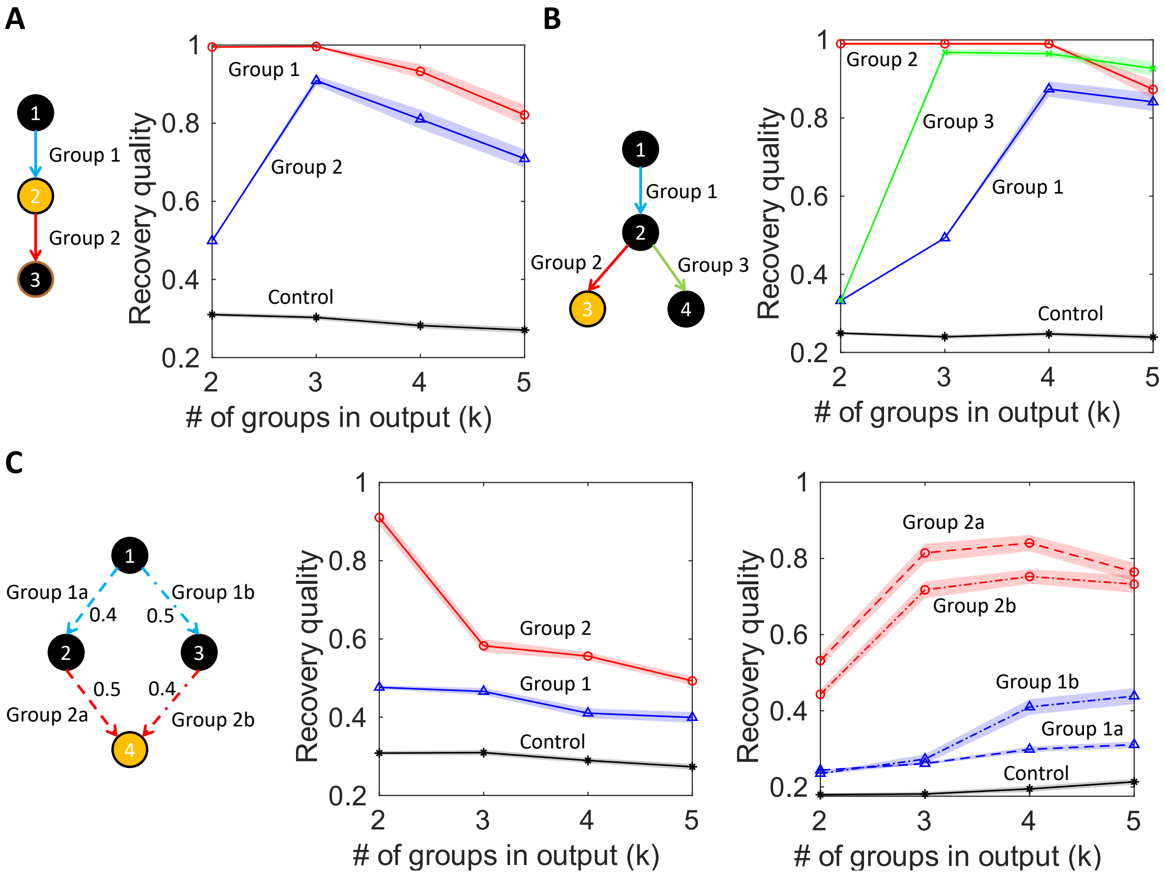


Figure S3: **Metropolis is able to identify groups associated with functions other than end product of linear digradation chain.** The recovery quantity of each functional group as function of number of groups in output ($k$) for function to be (A) intermediate product of linear degradation chain; (B) one of the end product in a degradation chain with a branch; (C) common end product of 2 linear degradation chain. Groups are indicated in the pictogram of each panel. Numbers in the pictogram of (C) indicate the transfer ratio $w_r$ of each reaction. In the first two cases (A & B), Metropolis can identify all the functional groups. While in the last, it can only recover the group of direct producers (also see text).

### 5 Other supplementary figures

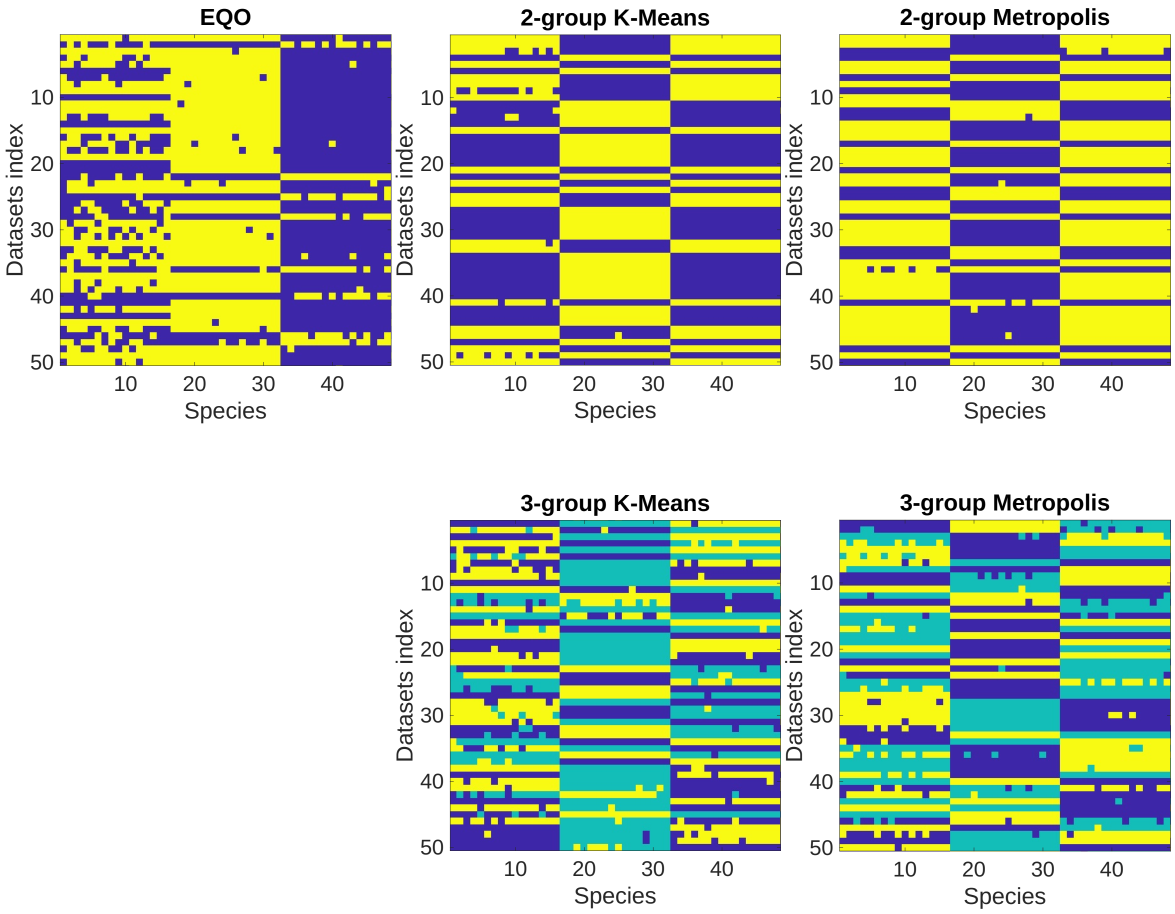


Figure S4: **Output groupings of the 3 algorithms for a linear degradation chain of $N = 3$ metabolites.** The output groupings of EQO, 2-group and 3-group K-means and Metropolis, as correspond to Fig. 2A&B in the main text. Species 1-16 belong to group 1, 17-32 belong to group 2 (direct producers), 33-48 belong to group 3 (nonfunctional species).

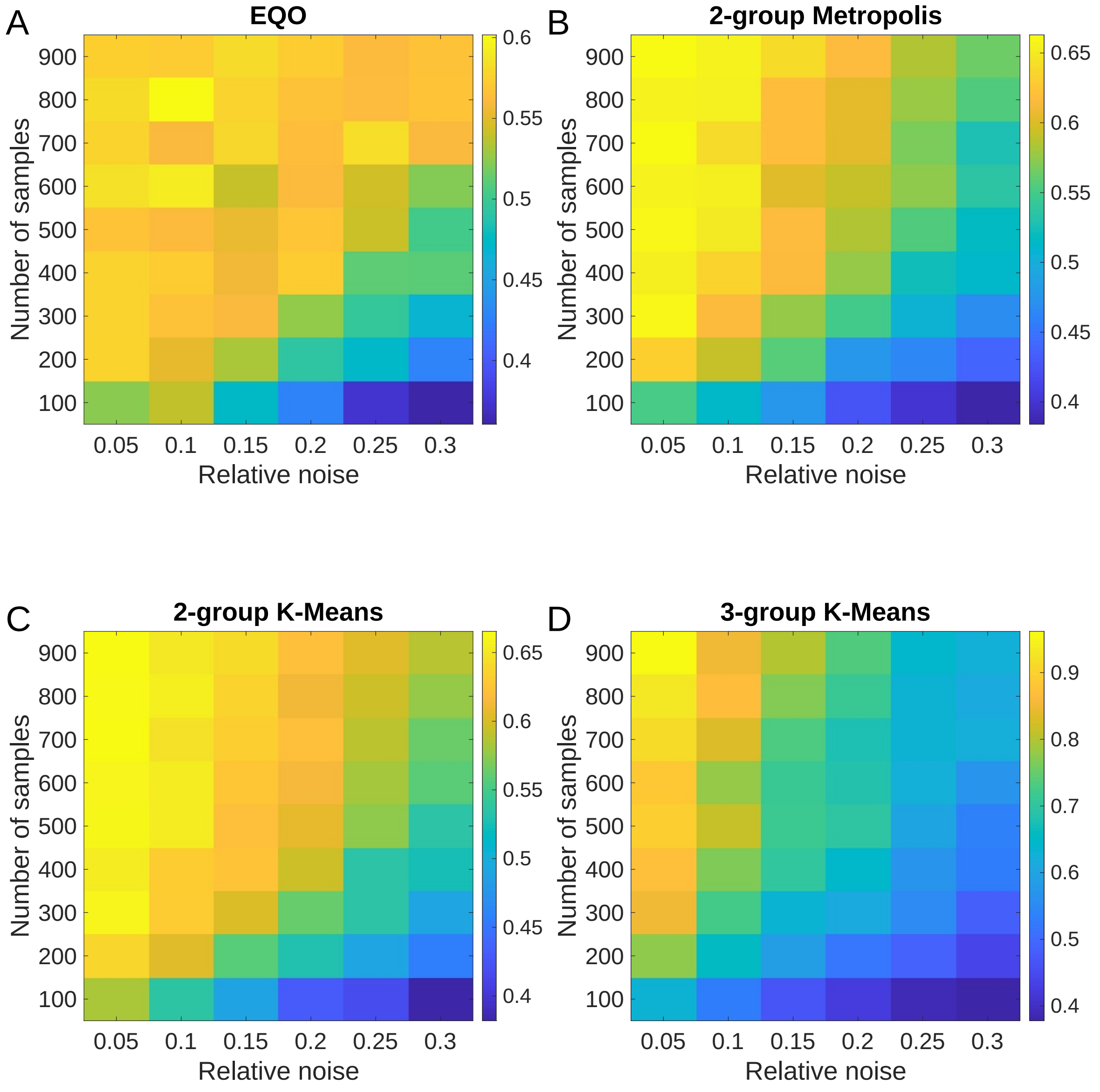


Figure S5: Heat maps of mean scores over 50 datasets of the algorithms as a function of relative noise and number of samples.
